# Gene flow and Andean uplift shape the diversification of *Gasteracantha cancriformis* (Araneae: Araneidae) in Northern South America

**DOI:** 10.1101/250977

**Authors:** Fabian C. Salgado-Roa, Carolina Pardo-Diaz, Eloisa Lasso De Paulis, Carlos F. Arias, Vera Nisaka Solferini, Camilo Salazar

## Abstract

**Aim:** The Andean uplift has played a major role shaping the current Neotropical biodiversity. However, in arthropods other than butterflies, little is known about how this geographic barrier has impacted species historical diversification. Here we examined the phylogeography of the widespread and color polymorphic spider *Gasteracantha cancriformis* to: (i) evaluate the effect of the northern Andean uplift on its divergence and, (ii) assess whether its diversification occurred in the presence of gene flow.

**Location:** Northern Andes and Brazil

**Methods:** We inferred phylogenetic relationships and divergence times in *G. cancriformis* using mitochondrial and nuclear data from 105 individuals in northern South America. Genetic diversity, divergence and population structure were quantified. We also compared multiple demographic scenarios for this species using a model-based approach (PHRAPL) to establish divergence with or without gene flow. Finally, we evaluated the association between genetic variation and color polymorphism.

**Results:** Both nuclear and mitochondrial data supported two well-differentiated clades, which correspond to populations occurring in opposite sides of the Eastern cordillera of the Colombian Andes. The splitting between these clades occurred in the early Pleistocene, around 2.13 million years ago (95% HPD = 0.98–3.93).

Despite this vicariant event, there is shared genetic variation between the clades, which is better explained by a scenario of historical divergence in the face of gene flow. Color polymorphism was randomly scattered in both clades and is not statistically associated with the genetic variation studied here.

**Main Conclusions:** The final uplift of Eastern cordillera of the Colombian Andes was identified as the major force that shaped the diversification of *G. cancriformis* in Northern South America, resulting in a *cis*- and *trans*-Andean phylogeographic structure for the species. The divergence in the face of gene flow between *cis*- and *trans*-Andean populations found for this spider has been likely facilitated by the presence of low-elevation passes across the Eastern Colombian cordillera. Our work constitutes the first example where the Andean uplift coupled with gene flow influenced the evolutionary history of an arachnid lineage.

## INTRODUCTION

The Northern Andes in South America is one of the most biodiverse regions on the planet, and the origins of this rich diversity has been linked to past geological and climatic events such as the uplift of the Andes and quaternary climatic oscillations (Kattan et al., 2004). The effect of these geoclimatic events in promoting divergence between Neotropical populations and species can be elucidated with genetic data, especially by detecting deviations from the expected coalescent patterns in neutral loci (Rull, 2011). Most studies addressing this question have identified the uplift of the Northern Andes as a major driver of Neotropical diversification in a scenario consistent with allopatric differentiation, wherein the complex topography of the Andes isolated populations on both sides of this barrier thus restricting gene flow (Antonelli et al., 2009; Brumfield & Capparella, 1996; Hoorn et al., 2010). In contrast with this classic view, a recent comparative phylogeographic study found discordant divergence times for multiple avian lineages with cross-Andean distribution, a result that is better explained by dispersal ability across the Andes rather than a single vicariant event (Smith et al., 2014). In line with this finding, new evidence supports the notion that common diversification modes in Neotropical birds include secondary contact between cross-Andean populations or divergence in presence of gene flow facilitated by low-elevation corridors along the Andes (Cadena, Pedraza, & Brumfield, 2016; Oswald et al., 2017).

However, our current knowledge on the modes of animal diversification in the Northern Andes is mostly based in vertebrates and, although arthropods are the most diverse group of animals, analyses of their diversification in this region remain scarce (De-silva et al., 2017; Turchetto-Zolet et al., 2013). Some studies limited to insects, especially butterflies, show that the Andean mountains have had an important role triggering their diversification, where speciation with and without gene flow across the Andes has occurred (Arias et al., 2014; Chazot et al., 2016, 2017; De-silva et al., 2017; Díaz et al., 2014; Dick, Roubik, Gruber, & Bermingham, 2004; Elias et al., 2009). Yet, a comprehensive understanding on how the Andean orogeny has promoted Neotropical animal diversification needs the inclusion of additional arthropod taxa, like arachnids.

*Gasteracantha cancriformis* (Linnaeus, 1758) is a Neotropical orb-web and color polymorphic spider that has at least eight known morphs (Gawryszewski, 2007). This color polymorphism, however, is still enigmatic (Gawryszewski & Motta, 2012). The species has a wide distribution in the Neotropical region, occurring in both sides of the Andes and in the Colombian inter-Andean valleys. This distribution makes it a great model to test whether the uplift of the Andes has influenced its diversification at the population level.

Here, we implemented a multilocus approach to study the genetic connectivity between polymorphic populations of *G. cancriformis* across the Northern Andes (Fig. 1, table S1) and tested scenarios of strict vicariance vs. diversification with gene flow. We also evaluated if lineage clustering in this spider is explained by geography or color pattern. Overall, this work contributes to deepen our understanding on how Andean orogeny has shaped processes of biotic diversification and biogeography in the Neotropics.

**Figure 1.**
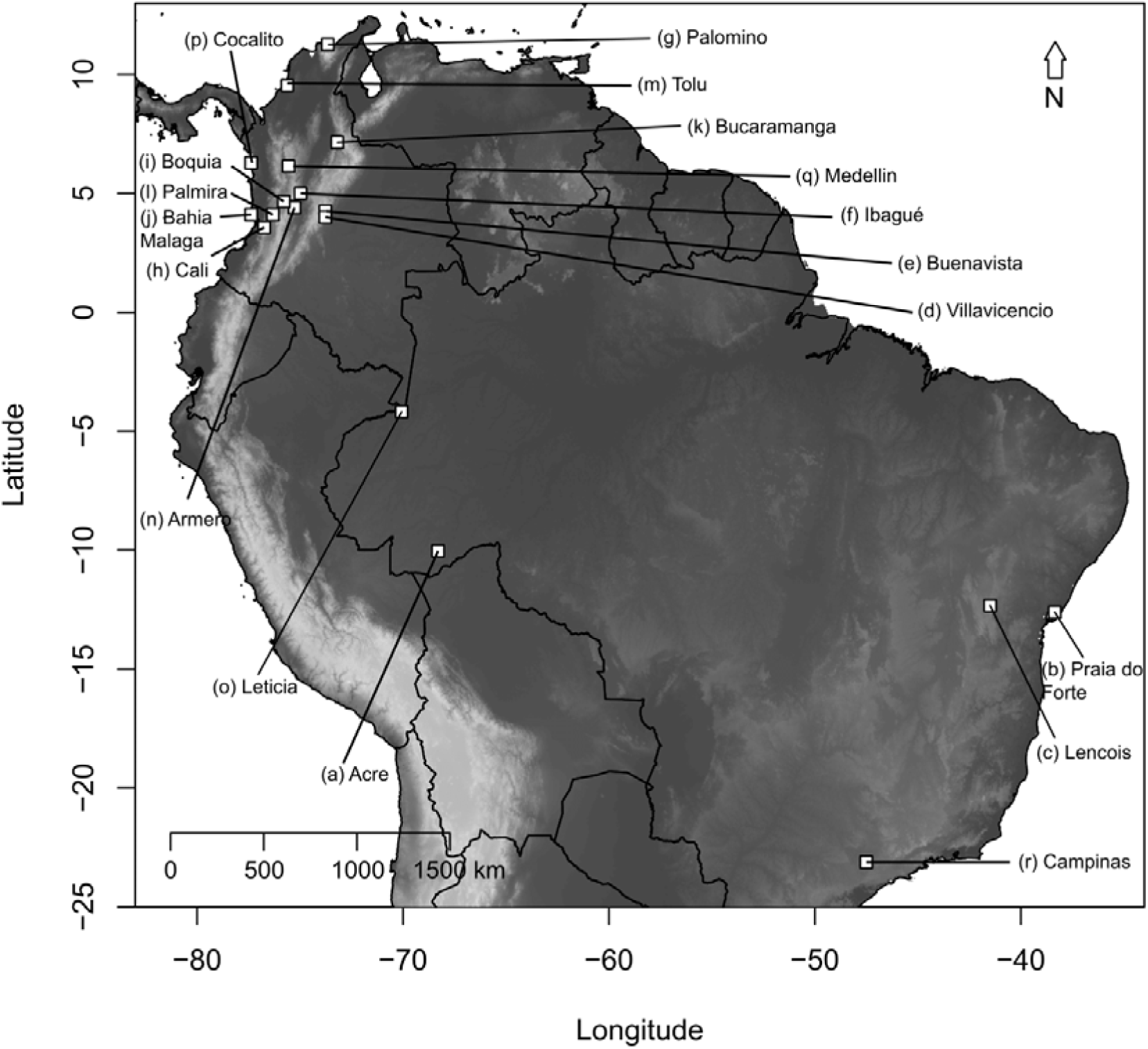
Map of the Neotropics showing the 18 sampling localities in Colombia and Brazil. White squares correspond to each locality sampled. Sampled localities are: (a) Acre, (b) Praia do forte, (c) Lencois, (d) Villavicencio, (e) Buenavista, (f) Ibague, (g) Palomino, (h) Cali, (i) Boquia, (j) Bahia Malaga, (k) Bucaramanga, (l) Palmira, (m) Tolu, (n) Armero, (o) Leticia, (p) Cocalito, (q) Medellin and (r) Campinas.

## MATERIALS AND METHODS

### Sample collection

We used standard aerial searching and beating methods to sample a total of 105 individuals of *G. cancriformis* from 17 localities distributed from the North of Colombia to Southeastern Brazil (Fig. 1, Table S1). Specimens were color coded following Gawryszewski (2007), preserved in a 20% dimethylsulphoxide (DMSO) solution saturated with NaCl and stored at −80 °C. Colombian samples were deposited in the Arthropods Collection of the Universidad del Rosario (CAUR#229) and Brazilian specimens were deposited in the Coleção Científica de Aracnídeos e Miriápodes of the Instituto Butantan (São Paulo, Brazil).

### DNA extraction, amplification and sequencing

DNA was extracted from legs using the Wizard Genomic DNA purification kit (Promega), following the manufacturer’s protocol. We amplified two mitochondrial genes, the Cytochrome Oxidase I (587 pb; COI; Folmer, Black, Hoeh, Lutz, & Vrijenhoek, 1994) and the 16S ribosomal DNA gene (397 pb; 16s; Simon et al., 1994), using the conditions reported elsewhere (Blackledge et al., 2009; McHugh, Yablonsky, Binford, & Agnarsson, 2014; Peres et al., 2015). We also sequenced two nuclear loci, the internal transcribed spacer 2 (296 pb; ITS; White, Bruns, Lee, & Taylor, 1990) and 28S ribosomal DNA gene (681 pb; 28s; Hedin & Maddison, 2001), which have been previously used in spiders (Moradmand, Schönhofer, & Jäger, 2014; Peres et al., 2015). Finally, we used the published transcriptome of *G. cancriformis* (Prosdocimi et al., 2011) and designed the primers forward: 5’- CAATTACACCTGGGAATCTTCTG-3’ and reverse: 5’- CCCTGACAAAATTCAAATAGTCG-3’ to amplify a 560 bp fragment of the heat shock protein 90 (HSP90), a gene that was used previously in phylogenetic studies in *Heliconius* (Salazar et al., 2010). PCR reactions for this marker were performed in a 10μL reaction volume containing 1× PCR buffer, 2.5 mM MgCl_2_, 500 μM dNTPs, 0.5μM each primer, 0.5 U Platinum Taq (INVITROGEN) and 30–40 ng of DNA. The PCR cycling profile was 94°C for 5 min, 40 cycles at 94°C for 30 s, Tm 54°C for 30 s and 72°C for 1 min and final extension at 72°C during 10 min. For all loci, we visualized 1μL of the PCR reaction in a 1.5% agarose gel to verify the success of PCR. Positive amplicons were cleaned up by ExoSAP-IT (USB Corp., Cleveland, OH) and their sequencing was carried out by ELIM Biopharmaceuticals Inc. (Hayward, CA).

Gene sequences were read, edited and assembled with CLC MAIN WORKBENCH (QIAGEN), to obtain a consensus sequence per-individual. For nuclear loci, haplotype inference for heterozygous calls was conducted with the PHASE algorithm implemented in DNASP V5.10 (Librado & Rozas, 2009) with 5,000 iterations per simulation and accepting inferred haplotypes with a confidence higher than 90%. Then, we used MEGA 6.0 (Tamura, Stecher, Peterson, Filipski, & Kumar, 2013) using the CLUSTAL W algorithm (Thompson, Higgins, & Gibson, 1994) to create alignments for each loci; these alignments were visually inspected and corrected. Alignments were translated to protein and verified for stop codons in MESQUITE V3.04 (Maddison & Maddison, 2015).

### Molecular phylogenetics and divergence times

The nucleotide substitution model for each mitochondrial gene was selected using the BIC criterion in jMODELTEST 0.1.1 (Posada, 2008). The most suitable models were HKY+I for COI and TIM+I for 16S.

Tree topologies were estimated with Bayesian inference (BI) using BEAST 1.7.4 (Drummond, Suchard, Xie, & Rambaut, 2012) and including two *Micrathena vigorsi* individuals as outgroups (Table S1). We unlinked the substitution model for each gene and linked the clock model and tree. We applied a lognormal relaxed clock to estimate divergence times using a mutation rate of 0.0112 (SD=0.001) substitution/site/million years, previously reported for node dating and calibration in spiders (Bartoleti et al., 2017; Bidegaray-Batista et al., 2011; Kuntner, Arnedo, Trontelj, Lokovšek, & Agnarsson, 2013). We ran two runs of 100 million generations sampling every 1000 generations. The initial 10000 trees were discarded as burn-in using TREEANNOTATOR (DRUMMOND & BOUCKAERT, 2015). We examined the output in TRACER1.5 (Rambaut & Drummond, 2014) to confirm that all effective sample size (ESS) values were greater than 200 and the convergence of the chains to the stationary distribution. The maximum credibility tree that best represented the posterior distribution was visualized and edited with FIGTREE1.4 (Rambaut, 2012).

Phylogenetic reconstruction was also done with Maximum Likelihood (ML) in IQ-TREE (Nguyen, Schmidt, Von Haeseler, & Minh, 2015) using the same substitution models described before and applying the edge-proportional partition. Node support was assessed with 1000 bootstrap replicates.

### Population genetics

We calculated haplotype (h) and nuclear (π) diversity, number of segregating sites (ss) and Tajima’s D with DNASP V5.10 (LIBRADO & ROZAS, 2009). Genetic structure was evaluated using F_ST_ at two different levels: (i) among phylogenetic clades (i.e. populations occurring in opposite sides of the Eastern cordillera of the Colombian Andes) and (ii) among populations, and significance of deviations from panmixia was assessed with the Hudson’s permutation test (Hudson, Boos, & Kaplan, 1992). An analysis of molecular variance (AMOVA) was also calculated for the same levels of differentiation with ARLEQUIN 3.5 (Excoffier & Lischer, 2010) using 10000 permutations.

Using the nuclear dataset we identified the number of population genetic clusters (K) with the Bayesian clustering approach implemented in STRUCTURE 2.3.4 (Pritchard, Stephens, & Donnelly, 2000). We ran the analysis under the admixture model, with a 50,000 burn-in and 100,000 MCMC sampling generations for K ranging from 1 to 13 (localities with only one individual were removed from this analysis), with 20 iterations for each value of K. We determined the K that better reflects our data applying three complementary approaches as recommended by Janes et al. (2017): (i) according to the delta K method of Evanno (Evanno, Regnaut, & Goudet, 2005), (ii) plotting the likelihood of K for each value of K (Earl & vonHoldt, 2012) and, (iii) reporting multiple barplots for K values between 2 and 5. All these tests were implemented in CLUMPAK (Kopelman et al., 2015). An additional validation of the genetic clusters for each locus was done with multivariate analysis. For this, fasta sequences were transformed into a genind object and loaded into the *adegenet* R package (Jombart & Ahmed, 2011), where we performed a principal component analysis (PCA). We retained the first two components for a subsequent Canonical Discriminant Analysis using the R package *candisc (Friendly & Fox, 2017)*.

As isolation by distance (IBD) can obscure population structure signals, we investigated the presence of IBD for each locus using Mantel (Mantel, 1967) with the R package *vegan (Dixon, 2003)*. For this, pairwise geographic distances among localities were calculated with the function *distm* from R package *geosphere* (Hijmans, 2016) while genetic distances were estimated by linearizing the F_ST_ values obtained previously. We also implemented a partial Mantel test (Smouse, Long, & Sokal, 1986) to separate the effect of geographic distance from the population assignments, based on STRUCTURE results.

Considering the recent concerns about Mantel test (Legendre, Fortin, & Borcard, 2015; Legendre & Fortin, 2010; Meirmans, 2012), we also tested linear correlations between the logarithm of the geographic distances and genetic distances as recommended by Legendre & Fortin (2010) and Diniz-Filho et al. (2013).

### Demographic model testing

We used PHRAPL (Jackson et al., 2017) to choose a demographic model that fits our data. PHRAPL compares the topologies obtained from empirical data with those simulated under multiple demographic models and then, by calculating the proportion of times that simulated topologies match the empirical ones, it approximates the log-likelihood of the data given a model. PHRAPL calculates Akaike Information Criterion (AIC) as a measure of lack of model fit, whilst the associated AIC weights (wAIC) can be interpreted as model probabilities.

PHRAPL can compare various demographic models, which can be broadly categorized as: (i) isolation-only (IO), (ii) migration-only (MO), (iii) isolation-with-migration (IM) and, (iv) mixed models (ME). We tested six models that fall into the IO and IM categories. The first is an IO model with a single coalescent event and no migration, while the other five are IM models that assume constant gene flow along the branch length and differ in the direction and strength of migration (Fig. 4).

As input for PHRAPL we assigned each individual to its collecting site (east or west of the Eastern cordillera) and loaded five midpoint rooted trees estimated with IQTREE (one per locus; Nguyen et al., 2015). We used jMODELTEST (Posada, 2008) to select the most plausible pattern of sequence evolution for each gene (COI: HKY+I, 16S: TIM+I, ITS: K80+I, 28S: F81+I, HSP90: GTR+I)

We subsampled four tips per group and 100 subsamples per gene, giving a total of 500 observed trees. Then, 100,000 trees were simulated for each model using a grid of parameter values for population divergence (t = 0.30,0.58,1.40,2.54,4.1) and migration (m = 0.10,0.22,0.46,1.00,2.15), in units of 4N and 4Nm respectively. In case gene flow was detected in our dataset, we tested two additional models that correspond to recent (τ-9τ/10) and ancient secondary contact (τ-τ/5), starting from the tips.

### Phenotype by genotype association

To test whether there is an association between the coloration of individuals and their genetic variation, we ran a chi-square Monte Carlo test under the null hypothesis of independence between coloration and genetic haplotypes. This analysis was run for each locus and establishing color morphs categories following Gawryszewski (2007). The analysis included only Colombian individuals as Brazilian samples do not have color records.

## RESULTS

### Phylogenetic relationships and divergence time

Both BI and ML showed the same phylogenetic pattern where two highly supported clades are recovered, with their internal relationships unresolved (Fig. 2). These major clades correspond to populations at the eastern (eEC) and western (wEC) sides of the eastern cordillera of the Colombian Andes (EC), with some individuals from the foothills of the EC showing shared haplotypes between both clades (Fig. 2). Other physiographic features in the Neotropics, such as the western and central cordilleras of the Colombian Andes or the Brazilian dry diagonal, do not cause population structure in *G. cancriformis*. Accordingly, Brazilian samples were monophyletic within the eEC clade (Fig. 2). The divergence time for the two main clades was estimated at *ca.* 2.13 Ma (95% HPD = 0.98–3.93 Ma), which is very close to the Pliocene/Pleistocene boundary and concordant with the final EC uplift (Gregory-Wodzicki, 2000).

**Figure 2.**
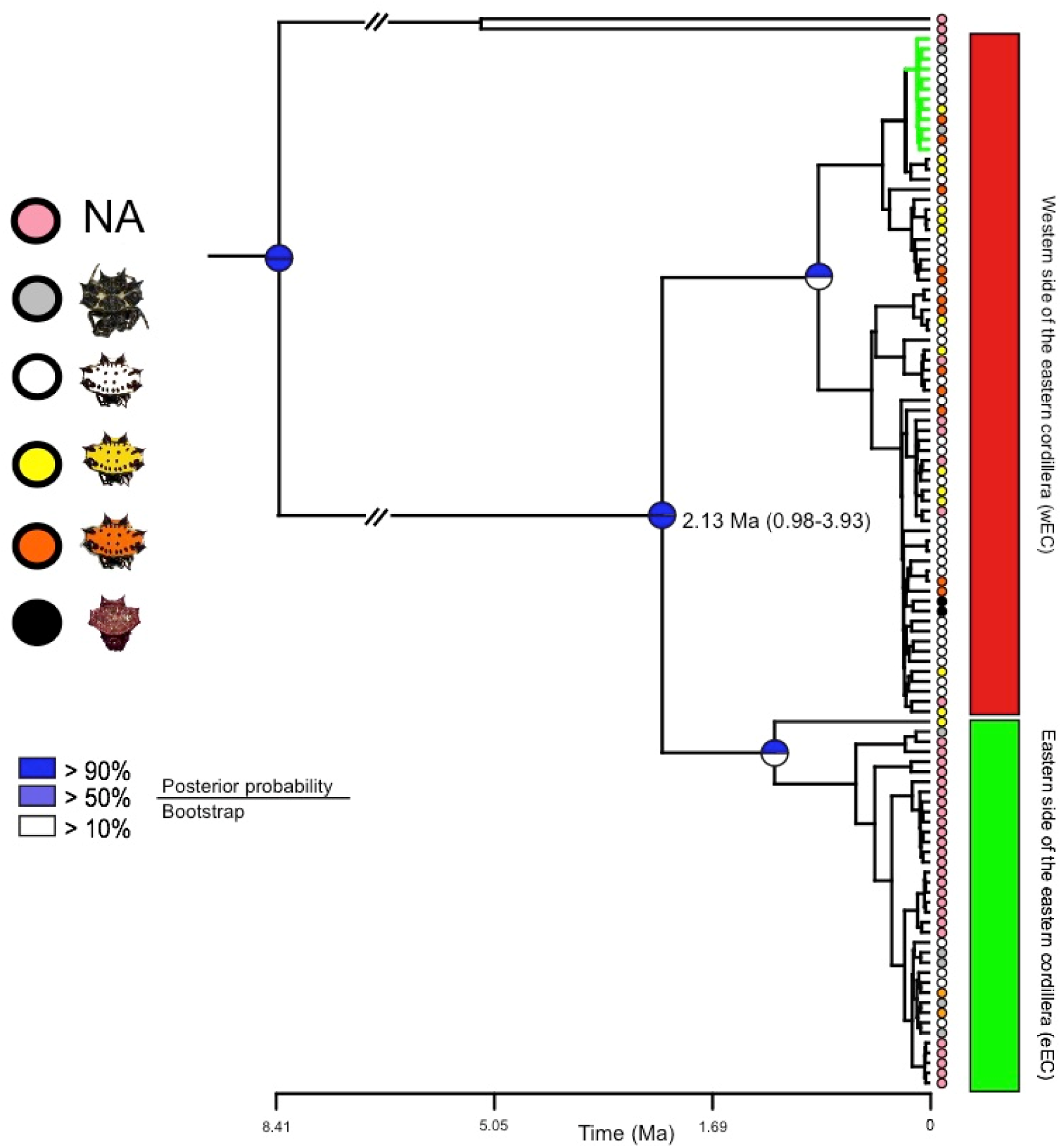
Mitochondrial phylogeny. Best recovered tree with mtDNA where node supports are represented by circles divided in two: the upper half corresponds to posterior probabilities obtained by Bayesian Inference and the lower half to the Maximum-Likelihood boostrap values after 1000 bootstrap pseudoreplicates. Colored circles at the tips represent the color phenotype in the opisthosoma of each individual. Green and red squares highlight the eastern (eEC) and western (wEC) sides of eastern cordillera of the Colombian Andes. Green branches highlight individuals sampled in the eEC that cluster into the wEC clade.

## Population Genetics

Mitochondrial and nuclear sequences were clustered in two genetically differentiated groups, corresponding to the eEC and wEC clades (F_ST__mtDNA=0.60; F_ST__ITS=0.24; F_ST__28S=0.25; F_ST__HSP90=0.20; for all loci P<0.05 in the Hudson’s permutation test). Mitochondrial nucleotide diversity was higher in the eEC clade than in the wEC clade (Table 1); however, this did not hold true for nuclear loci, which may be due to differences in effective size and other causes of mito-nuclear discordance (Toews & Brelsford, 2012). None of the loci showed significant Tajimas’ D, suggesting neutral evolution (Table 1). For all loci, genetic structure was more pronounced among populations sampled at different sides of the EC than among populations at the same side (Fig. S1). This pattern was also reflected in the AMOVA analysis where part of the variation is explained by differences among regions (eEC and wEC clades; Table S2), although for nuclear loci, most of the variance is due to differences within population.

**Table 1.**
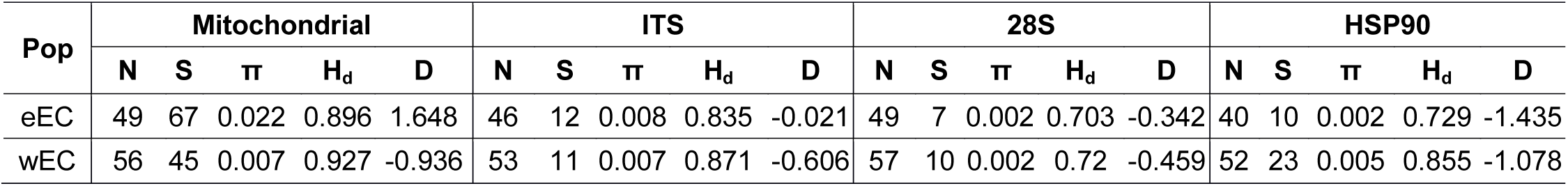
Population genetic summary statics for the eastern (eEC) and western (wEC) sides of the Eastern Colombian cordillera for each loci. Population (Pop), number of sequences (N), number of polymorphic sites (S), nucleotide diversity (π), haplotype diversity (Hd) and Tajima’s D (D). *P < 0.05; **P < 0.02; none of the loci showed Tajima’s D values that departed from neutral expectations.

All methods applied to select the optimal value of K consistently revealed two groups (K=2; Fig. 3, Fig. S2 & S3), which agrees with the phylogenetic analyses and the population pairwise F_ST_ values (Fig. 2 & Fig. S1). In agreement with those results, the canonical discriminant analyses also identified two geographical clusters (Fig. S4). Individuals from both groups share variation among them. For example, most individuals from Villavicencio (eastern foothills of the EC) presented either wEC or eEC mtDNA, whilst their nDNA was up to 30% from the wEC clade; even more, there were two individuals from this locality with wEC mtDNA and almost ~80% of their nDNA was of wEC origin (Fig. 2, Fig. 3 & Fig. S5). Likewise, two individuals from Boquia and Bucaramanga (west and central cordillera, respectively) presented wEC mtDNA but their nDNA showed almost 50% of shared ancestry with the eEC populations (Fig. 2, Fig. 3 & Fig. S5). We ruled out any effect of isolation by distance rather than Andean divergence causing the geographical population structure observed here (Fig. S6, Table S3).

**Figure 3.**
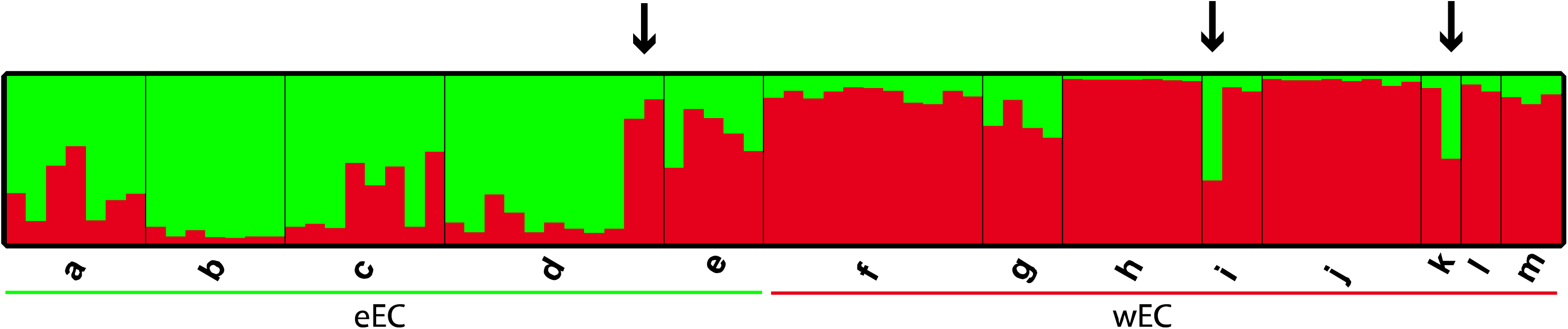
Bayesian population assignment test based on nDNA. A population assignment test with the software STRUCTURE based on three nuclear loci identified two distinct populations (K=2). Bar plots show Bayesian assignment probabilities for individuals where each color represents the most likely ancestry from which the genotype was derived (green: eEC and red: wEC). Bars on the bottom indicate the geographical region each population belongs to. Populations are coded as in Figure 1. In population d, individuals 67 and 68 (arrow), have almost ~80% of their nDNA from wEC. Individual 78 in population i (arrow) and individual 95 in population m (arrow) have wEC mtDNA but their nDNA showed almost 50% of shared ancestry with the eEC populations.

**Figure 4.**
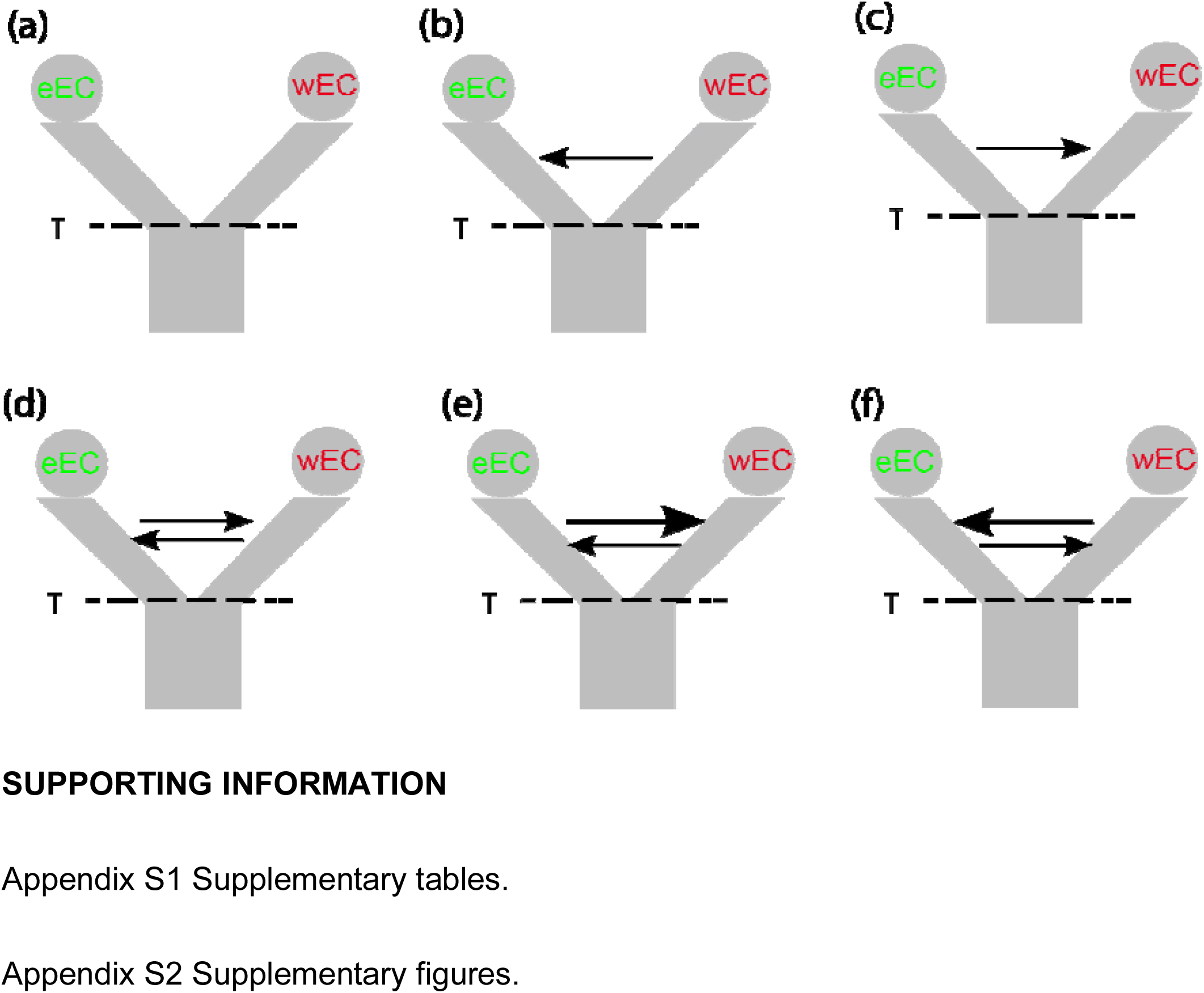
Demographic scenarios tested for the evolution of *G. cancriformis* in the Northern Andes. a) Divergence with no migration, b) Divergence with unidirectional migration from wEC to eEC, c) Divergence with unidirectional migration from eEC to wEC, d) Divergence with bidirectional symmetrical migration, e) Divergence with bidirectional asymmetrical migration from eEC to wEC and, f) Divergence with bidirectional asymmetrical migration from wEC to eEC.

### Demographic models

PHRAPL revealed different wAIC values and high ∆AIC between models including gene flow (IM) and isolation only (IO; Table 2 & Table S4). The model with no migration had the lowest wAIC, indicating that a single vicariant event with no genetic exchange is not plausible yet, it is difficult to differentiate between symmetrical vs. asymmetrical gene flow (Table 2 & Table S4). When we tested recent vs. ancient secondary contact, the latter model was better supported, ruling out recent secondary contact but suggesting at least some isolation caused by the vicariant event (Table 2). These results imply that gene flow is responsible for the *G.cancriformis* shared ancestry between eEC and wEC geographical regions.

**Table 2.**
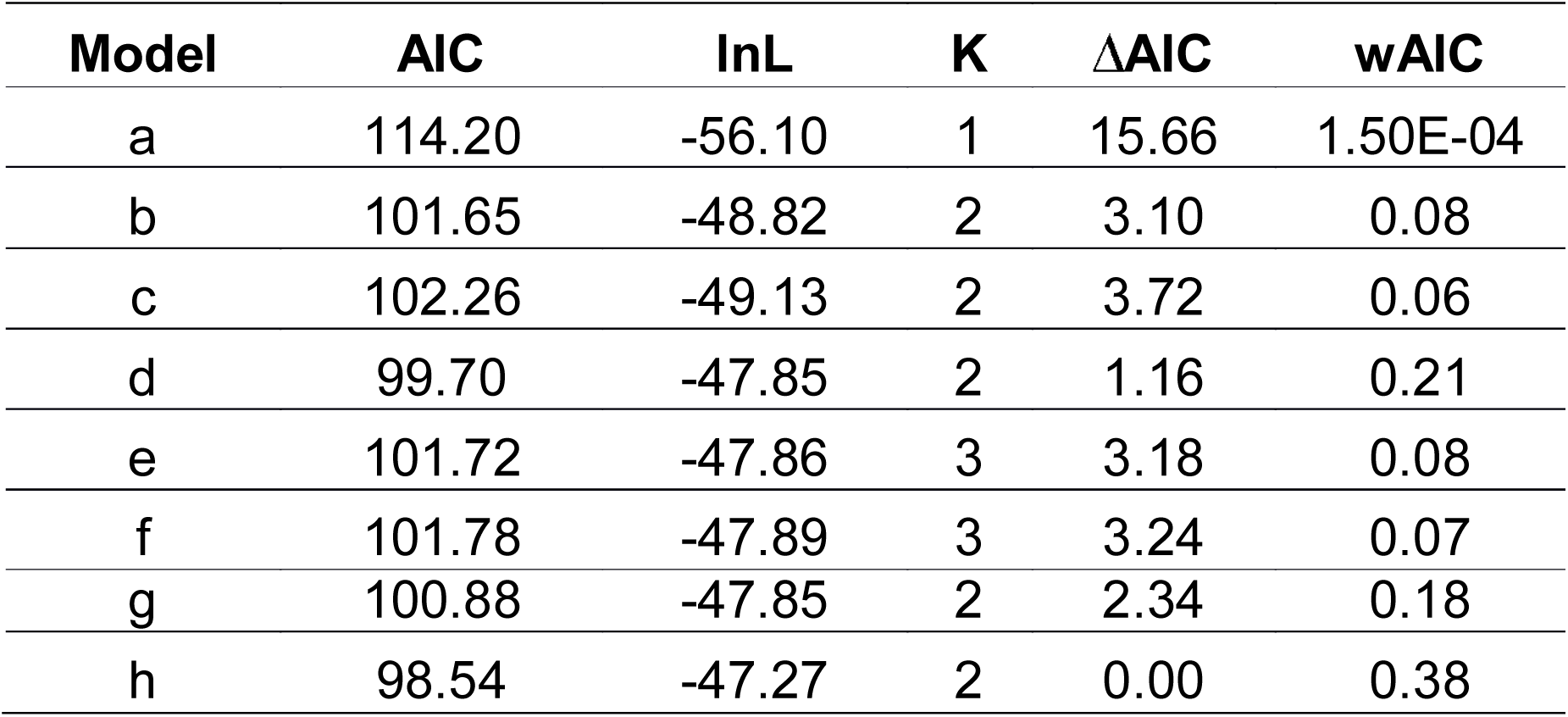
Measures of fit of alternative models, labeled as in Fig. 4. Isolation only (a), Isolation with Migration (b to f) and Secondary contact (g and h).

| **Model** | **AIC** | **lnL** | **K** | **ΔAIC** | **wAIC** |
| --- | --- | --- | --- | --- | --- |
| a | 114.20 | -56.10 | 1 | 15.66 | 1.50E-04 |
| b | 101.65 | -48.82 | 2 | 3.10 | 0.08 |
| c | 102.26 | -49.13 | 2 | 3.72 | 0.06 |
| d | 99.70 | -47.85 | 2 | 1.16 | 0.21 |
| e | 101.72 | -47.86 | 3 | 3.18 | 0.08 |
| f | 101.78 | -47.89 | 3 | 3.24 | 0.07 |
| g | 100.88 | -47.85 | 2 | 2.34 | 0.18 |
| h | 98.54 | -47.27 | 2 | 0.00 | 0.38 |

### Phenotype by genotype association

We found the white phenotype as the most frequent morph in all populations, while the black-white morphology was only present in the eEC populations. Nonetheless, in the Colombian Cauca valley (wEC), we collected a black morph that has not been previously reported (Fig. S7). However, our molecular sampling (mtDNA and nDNA) revealed a statistical association between genetic variation and geography, but such association was not found for color polymorphism (Table S5). This is also evident in the mtDNA phylogeny, where individuals of different colors group in the same clade (Fig. 2).

## DISCUSSION

Our mitochondrial and nuclear data consistently showed two well-supported genetic clusters separated by the EC of the Colombian Andes. The divergence of these *G. cancriformis* groups occurred during the late Miocene and early Pleistocene, which is coincident with the final uplift of this part of the Andes (Gregory-Wodzicki, 2000). In contrast, the western and central Colombian cordilleras as well as the dry diagonal in Brazil do not seem to contribute to the diversification of these spider populations. This contrasts with previous reports where the Brazilian dry diagonal has been found as a natural barrier to gene flow in taxa such as frogs, birds and lizards (Fouquet et al., 2014; Harvey & Brumfield, 2015; Werneck et al., 2012). Also, the genetic homogeneity found within the wEC clade may be explained by the topography itself of the western and central Colombian cordilleras, which are considerably narrower than the eastern one (Guarnizo et al., 2015), likely facilitating dispersal of individuals across this region.

Species diversification driven by the uplift of the Andes has been documented in several organisms including birds (Chaves, Pollinger, Smith, & LeBuhn, 2007; Fernandes, Wink, Sardelli, & Aleixo, 2014; Ribas, Moyle, Miyaki, & Cracraft, 2007), reptiles (Teixeira et al., 2016), amphibians (García-R et al., 2012; Guarnizo et al., 2015; Guarnizo, Amézquita, & Bermingham, 2009), bees (Dick et al., 2004) and butterflies (Elias et al., 2009). Yet, the persistence of gene flow between populations separated by the Andes is far less known (Hoffmann & Baker, 2003; Miller et al., 2008; Oswald et al., 2017). Here, despite the vicariance associated with the Andean uplift that resulted in eEC and wEC Andean clades for *G. cancriformis*, we found individuals with shared ancestry between the main two geographical groups. The approximate likelihood demographic model implemented identified gene flow as the most likely explanation for this. Furthermore, the model with the best support implies divergence in the face of gene flow after τ/5 generations forward in time, which suggests a short allopatric period.

Altitudinal depressions across the Andes can contribute to dispersal of individuals, thus allowing admixture between populations that occur at opposite sides of this barrier. In fact, the EC of the Colombian Andes is not a uniform barrier along its length and counts with at least two depressions, the Andalucía pass and the Suaza-Pescado valleys, which may be acting as dispersal corridors (Cadena et al., 2016). To our knowledge, scenarios consistent with Andean altitudinal depressions facilitating dispersion and gene flow have only been reported in the Peruvian Porculla pass, where six co-distributed bird taxon pairs showed asynchronous divergence times likely due to independent dispersal events coupled with gene flow (Oswald et al., 2017). In arthropods, there is evidence for dispersal through the Andes (Dick et al., 2004), but the persistence of gene flow across this barrier has not been shown. We hypothesize that eEC and wEC populations of *G. cancriformis* have used such kind of passes to cross the Eastern Cordillera and reproduce with populations in opposite sides, even after they have achieved some degree of divergence. This could be facilitated by aerial dispersal mechanisms like ballooning, where the friction between air and the spider with its silk can make an individual move up to 3,200 Km (Gressitt, 1965). Although this displacement strategy has not been observed in *G. cancriformis*, it is used by its sister taxa (Bell et al., 2005).

Color polymorphism in the opisthosoma of *G. cancriformis* did not explain the structure found in this species. However, the lack of association of mtDNA or nDNA haplotypes with coloration may be due to the nature of the loci studied, as they evolve neutrally and are not members of any known pigmentation pathway in arthropods (Wittkopp & Beldade, 2009). Even so, the mtDNA phylogenetic pattern suggests, to some extent, that this polymorphism pre-dates the geographical split. Alternatively, the genetic connectivity between the populations at both sides of the Andes may be favoring the flow of color alleles thus maintaining phenotypic variation. This remains to be tested.

The scenario of vicariance coupled with gene flow found here support the original ideas of Chapman (1917, 1926) and Haffer (1967a, 1967b), who claimed that the similarities in the composition of the flora and fauna at both sides of the Andes might be due to dispersals through altitudinal depressions in the cordilleras (Chapman, 1917, 1926; Haffer, 1967, 1967).This case constitutes one of the few phylogeographic studies in arthropods, and the first arachnids showing that the Andes act as a permeable barrier, rather than an absolute one, for the free movement of alleles.

## Data availability

Sequences were deposited in GenBank under accession numbers xxxx-xxxx.

## Author contributions

F.C.S-R., E.L.P., C.S. and V.N.S. conceived the idea and designed the experiments. F.C.S.-R. collected the individuals. F.C.S.-R. and C.P-D. processed the samples in the laboratory and got the sequences. V.N.S. C.S. and C.F.A. contributed with material, tools and reagents. F.C.S-R. and C.S. analyzed the data. F.C.S.-R, C.S. and C.P-D wrote the manuscript. All authors approved the final version submitted.

## SUPPORTING INFORMATION

Appendix S1 Supplementary tables.

Appendix S2 Supplementary figures.

## ACKNOWLEDGEMENTS

For collecting permits, we thank Ministerio de Ambiente y Desarrollo Sostenible and ANLA in Colombia (Permiso Marco # 0530) and ICMBio in Brazil (27127 and 38889). We also thank multiple volunteers and friends for their logistic support and help with fieldwork in Colombia and Brazil. We are very grateful to Ariadna Morales for helping with the implementation of PHRAPL. FCS was funded by COLCIENCIAS (Joven Investigador Program Call 761-2016, contract FCN1705-CE101). CS and CPD were funded by COLCIENCIAS Grant FP44842-005-2017. CFA was supported by Convocatoria ‘Es tiempo de volver’-COLCIENCIAS-2014, contract No. 656–2014. NVS was funded by FAPESP (grants #2012/ 02526-7, #2013/50491-0 and #2013/08293-7). All authors declare no conflict of interest.

## REFERENCES

Antonelli A., Quijada-Mascareñas A., Crawford A. J., Bates J. M., Velazco P. M., & Wüster W. (2009). Molecular Studies and Phylogeography of Amazonian Tetrapods and their Relation to Geological and Climatic Models. In Amazonia: Landscape and Species Evolution (pp. 386–404). Wiley-Blackwell Publishing Ltd. http://doi.org/10.1002/9781444306408.ch24

Arias C. F., Salazar C., Rosales C., Kronforst M. R., Linares M., Bermingham E., & McMillan W. O. (2014). Phylogeography of Heliconius cydno and its closest relatives: Disentangling their origin and diversification. Molecular Ecology, 23(16), 4137–4152. http://doi.org/10.1111/mec.12844

Bartoleti, L. F. de M., Peres E. A., Sobral-Souza T., Fontes F., von H. M., Silva M. J. da, & Solferini V. N. (2017). Phylogeography of the dry vegetation endemic species Nephila sexpunctata (Araneae: Araneidae) suggests recent expansion of the Neotropical Dry Diagonal. Journal of Biogeography, 44(9), 2007–2020. http://doi.org/10.1111/jbi.12998

Bell J. R., Bohan D. A., Shaw E. M., & Weyman G. S. (2005). Ballooning dispersal using silk: world fauna, phylogenies, genetics and models. Bulletin of Entomological Research, 95(02), 69–114. http://doi.org/10.1079/BER2004350

Bidegaray-Batista L., Arnedo M. A., Phillips M., Rambaut A., Liebowitz T., Chan L., & Wilson A. (2011). Gone with the plate: the opening of the Western Mediterranean basin drove the diversification of ground-dweller spiders. BMC Evolutionary Biology, 11(1), 317. http://doi.org/10.1186/1471-2148-11-317

Black M., Hoeh W., Lutz, R., & Vrijenhoek, R. (1994). DNA primers for amplification of mitochondrial cytochrome c oxidase subunit I from diverse metazoan invertebrates, 3, 294–299.

Blackledge T. A., Scharff N., Coddington J. A., Szüts T., Wenzel J. W., Hayashi C. Y., & Agnarsson I. (2009). Reconstructing web evolution and spider diversification in the molecular era. Proceedings of the National Academy of Sciences of the United States of America, 106(13), 5229–34. http://doi.org/10.1073/pnas.0901377106

Brumfield R. T., & Capparella A. P. (1996). Historical Diversification of Birds in Northwestern South America: A Molecular Perspective on the Role of Vicariant Events. Evolution, 50(4), 1607. http://doi.org/10.2307/2410897

Cadena C. D., Pedraza C. A., & Brumfield R. T. (2016). Climate, Habitat Associations and the Potential Distributions of Neotropical Birds: Implications for Diversification Across the Andes. Rev. Acad. Colomb. Cienc. Ex. Fis. Nat., 40(155), 275–287.

Chapman F. M. (1917). The Distribution of Bird-Life in Colombia: A Contribution to a Biological Survey of South America.

Chapman F. M. (1926). The Distribution of Bird-Life in Ecuador: A Contribution to a Study of the Origin of Andean Bird-Life. Bulletin of the American Museum of Natural History, 55, 784.

Chaves J. A., Pollinger J. P., Smith T. B., & LeBuhn G. (2007). The role of geography and ecology in shaping the phylogeography of the speckled hummingbird (Adelomyia melanogenys) in Ecuador. Molecular Phylogenetics and Evolution, 43(3), 795–807. http://doi.org/https://doi.org/10.1016/j.ympev.2006.11.006

Chazot N., Willmott K. R., Condamine F. L., De-Silva D. L., Freitas A. V. L., Lamas G., & Elias M. (2016). Into the Andes: multiple independent colonizations drive montane diversity in the Neotropical clearwing butterflies Godyridina. Molecular Ecology, 25(22), 5765–5784. http://doi.org/10.1111/mec.13773

Chazot N., Willmott K. R., Lamas G., Freitas A. V. L., Piron-prunier F., Arias C. F., & Elias M. (2017). Renewed diversification following Miocene landscape turnover in a Neotropical butterfly radiation. bioRxiv.

De-silva D. L., Chazot N., Silva- K. L., Lamas G., Mallet J., Willmott K. R., & Elias M. (2017). North Andean origin and diversification of the largest ithomiine butterfly genus, 1–17. http://doi.org/10.1038/srep45966

Díaz S., Panzera F., Jaramillo-O N., Pérez R., Fernández R., Vallejo G., & Gómez-Palacio A. (2014). Genetic, Cytogenetic and Morphological Trends inthe Evolution of the Rhodnius (Triatominae: Rhodniini) Trans-Andean Group. PLoS ONE, 9(2), e87493. http://doi.org/10.1371/journal.pone.0087493

Dick C. W., Roubik D. W., Gruber K. F., & Bermingham E. (2004). Long-distance gene flow and cross-Andean dispersal of lowland rainforest bees (Apidae: Euglossini) revealed by comparative mitochondrial DNA phylogeography. Molecular Ecology, 13(12), 3775–3785. http://doi.org/10.1111/j.1365-294X.2004.02374.x

Diniz-filho J. A. F., Soares T. N., Lima J. S., Dobrovolski R., Landeiro V. L., Pires M., & Bini L. M. (2013). Mantel test in population genetics, 485, 475–485.

Dixon P. (2003). VEGAN, a package of R functions for community ecology. Journal of Vegetation Science, 14(6), 927–930. http://doi.org/10.1111/j.1654-1103.2003.tb02228.x

Drummond A. J., & Bouckaert R. R. (2015). Bayesian Evolutionary Analysis with BEAST. Cambridge: Cambridge University Press.

Drummond A. J., Suchard M. A., Xie D., & Rambaut A. (2012). Bayesian Phylogenetics with BEAUti and the BEAST 1.7. Molecular Biology and Evolution, 29 (8), 1969–1973. http://doi.org/10.1093/molbev/mss075

Earl D. A., & vonHoldt B. M. (2012). STRUCTURE HARVESTER: a website and program for visualizing STRUCTURE output and implementing the Evanno method. Conservation Genetics Resources, 4(2), 359–361. http://doi.org/10.1007/s12686-011-9548-7

Elias M., Joron M., Willmott K., Silva-BrandÃo K. L., Kaiser V., Arias C. F., & Jiggins C. D. (2009). Out of the Andes: Patterns of diversification in clearwing butterflies. Molecular Ecology, 18(8), 1716–1729. http://doi.org/10.1111/j.1365-294X.2009.04149.x

Evanno G., Regnaut S., & Goudet J. (2005). Detecting the number of clusters of individuals using the software structure: a simulation study. Molecular Ecology, 14(8), 2611–2620. http://doi.org/10.1111/j.1365-294X.2005.02553.x

Excoffier L., & Lischer H. E. L. (2010). Arlequin suite ver 3.5: a new series of programs to perform population genetics analyses under Linux and Windows. Molecular Ecology Resources, 10(3), 564–7. http://doi.org/10.1111/j.1755-0998.2010.02847.x

Fernandes A. M., Wink M., Sardelli C. H., & Aleixo A. (2014). Multiple speciation across the Andes and throughout Amazonia: the case of the spot-backed antbird species complex (Hylophylax naevius / Hylophylax naevioides), 1094–1104. http://doi.org/10.1111/jbi.12277

Fouquet A., Santana Cassini C., Fernando Baptista Haddad, C., Pech N., & Trefaut Rodrigues M. (2014). Species delimitation, patterns of diversification and historical biogeography of the Neotropical frog genus Adenomera (Anura, Leptodactylidae). Journal of Biogeography, 41(5), 855–870. http://doi.org/10.1111/jbi.12250

Friendly M., & Fox J. (2017). Visualizing Generalized Canonical Discriminant and Canonical Correlation Analysis. Vienna, Austria. Retrieved from https://cran.r-project.org/web/packages/candisc/candisc.pdf

García-R J. C., Crawford A. J., Mendoza, Á. M., Ospina O., Cardenas H., & Castro F. (2012). Comparative Phylogeography of Direct-Developing Frogs (Anura: Craugastoridae: Pristimantis) in the Southern Andes of Colombia. PLoS ONE, 7(9), e46077. http://doi.org/10.1371/journal.pone.0046077

Gawryszewski F. M. (2007). Policromatismo e stabilimentum em Gasteracantha cancriformis (Araneae, Araneidae): caracterização e as hipóteses da atração de presas e da proteção da teia. Universidade de Brasilia.

Gawryszewski F. M., & Motta P. C. (2012). Colouration of the orb-web spider Gasteracantha cancriformis does not increase its foraging success. Ethology Ecology & Evolution, 24(1), 23–38. http://doi.org/10.1080/03949370.2011.582044

Gregory-Wodzicki K. M. (2000). Uplift history of the Central and Northern Andes: A review. Geological Society of America Bulletin, 112(7), 1091–1105. http://doi.org/10.1130/0016-7606

Gressitt J. L. (1965). Biogeography and ecology of land arthropods of Antarctica., Biogeography and Ecology of Antarctica. Monographiae Biologicae, vol. 15 (pp. 431–490). Springer, Dordrecht.

Guarnizo C. E., Amézquita A., & Bermingham E. (2009). The relative roles of vicariance versus elevational gradients in the genetic differentiation of the high Andean tree frog, Dendropsophus labialis. Molecular Phylogenetics and Evolution, 50(1), 84–92. http://doi.org/https://doi.org/10.1016/j.ympev.2008.10.005

Guarnizo C. E., Paz A., Muñoz-Ortiz A., Flechas S. V., Méndez-Narváez J., & Crawford A. J. (2015). DNA Barcoding Survey of Anurans across the Eastern Cordillera of Colombia and the Impact of the Andes on Cryptic Diversity. PLOS ONE, 10(5), e0127312. http://doi.org/10.1371/journal.pone.0127312

Haffer J. (1967). Speciation in Colombian forest birds west of Andes. Novitates Zoologicae, 2294(2294), 1–57.

Haffer J. (1967). Zoogeographical Notes on the “Non-Forest” Lowland Bird Faunas of Northwestern South America. El Hornero, 10(4), 315–333.

Harvey M. G., & Brumfield R. T. (2015). Genomic variation in a widespread Neotropical bird (Xenops minutus) reveals divergence, population expansion, and gene flow. Molecular Phylogenetics and Evolution, 83, 305–16. http://doi.org/10.1016/j.ympev.2014.10.023

Hedin M. C., & Maddison W. P. (2001). A Combined Molecular Approach to Phylogeny of the Jumping Spider Subfamily Dendryphantinae (Araneae: Salticidae). Molecular Phylogenetics and Evolution, 18(3), 386–403. http://doi.org/10.1006/mpev.2000.0883

Hijmans R. J. (2016). geosphere: Spherical Trigonometry.

Hoffmann F. G., & Baker R. J. (2003). Comparative phylogeography of short-tailed bats (Carollia: Phyllostomidae). Molecular Ecology, 12(12), 3403–3414. http://doi.org/10.1046/j.1365-294X.2003.02009.x

Hoorn C., Wesselingh F. P., ter Steege H., Bermudez M. a, Mora, a, Sevink, J., & Antonelli, a. (2010). Amazonia through time: Andean uplift, climate change, landscape evolution, and biodiversity. Science (New York, N.Y.), 330(6006), 927–931. http://doi.org/10.1126/science.1194585

Hudson R. R., Boos D. D., & Kaplan N. L. (1992). A statistical test for detecting geographic subdivision. Molecular Biology and Evolution, 9(1), 138–151.

Jackson N. D., Morales A. E., Carstens B. C., & O’Meara B. C. (2017). PHRAPL: Phylogeographic Inference Using Approximate Likelihoods. Systematic Biology, 19, 431–435. http://doi.org/10.1093/sysbio/syx001

Janes J. K., Miller J. M., Dupuis J. R., Malenfant R. M., Gorrell J. C., Cullingham C. I., & Andrew R. L. (2017). The K = 2 conundrum. Molecular Ecology, 26(14), 3594–3602. http://doi.org/10.1111/mec.14187

Jombart T., & Ahmed I. (2011). adegenet 1. 3-1: new tools for the analysis of genome-wide SNP data, 27(21), 3070–3071. http://doi.org/10.1093/bioinformatics/btr521

Kattan G. H., Franco P., & Rojas V. (2004). Biological diversification in a complex region: a spatial analysis of faunistic diversity and biogeography of the Andes of Colombia, 1829–1839.

Kopelman N. M., Mayzel J., Jakobsson M., Rosenberg N. A., & Mayrose I. (2015). CLUMPAK: a program for identifying clustering modes and packaging population structure inferences across K. Molecular Ecology. http://doi.org/10.1111/1755-0998.12387

Kuntner M., Arnedo M. A., Trontelj P., Lokovšek T., & Agnarsson I. (2013). A molecular phylogeny of nephilid spiders: Evolutionary history of a model lineage. Molecular Phylogenetics and Evolution, 69(3), 961–979. http://doi.org/10.1016/j.ympev.2013.06.008

Legendre P., & Fortin M.-J. (2010). Comparison of the Mantel test and alternative approaches for detecting complex multivariate relationships in the spatial analysis of genetic data. Molecular Ecology Resources, 10(5), 831–844. http://doi.org/10.1111/j.1755-0998.2010.02866.x

Legendre P., Fortin M.-J., & Borcard D. (2015). Should the Mantel test be used in spatial analysis?Methods in Ecology and Evolution, 6(11), 1239–1247. http://doi.org/10.1111/2041-210X.12425

Librado P., & Rozas J. (2009). DnaSP v5: a software for comprehensive analysis of DNA polymorphism data, 25(11), 1451–1452. http://doi.org/10.1093/bioinformatics/btp187

Maddison W., & Maddison D. (2015). Mesquite: A modular system for evolutionary analysis, version 3.04. 2015 Available: http://mesquiteproject.org/mesquite/download/download.html.

Mantel N. (1967). The Detection of Disease Clustering and a Generalized Regression Approach. Cancer Research, 27(2 Part 1), 209 LP – 220.

McHugh A., Yablonsky C., Binford G. J., & Agnarsson I. (2014). Molecular phylogenetics of Caribbean Micrathena (Araneae: Araneidae) suggests multiple colonization events and single island endemism. Invertebrate Systematics, Accepted F(2012), 337–349. http://doi.org/10.1071/IS13051

Meirmans P. G. (2012). The trouble with isolation by distance. Molecular Ecology, 21(12), 2839–2846. http://doi.org/10.1111/j.1365-294X.2012.05578.x

Miller M. J., Bermingham E., Klicka J., Escalante P., do Amaral F. S. R., Weir J. T., & Winker K. (2008). Out of Amazonia again and again: episodic crossing of the Andes promotes diversification in a lowland forest flycatcher. Proceedings. Biological Sciences, 275(1639), 1133–42. http://doi.org/10.1098/rspb.2008.0015

Moradmand M., Schönhofer A. L., & Jäger P. (2014). Molecular phylogeny of the spider family Sparassidae with focus on the genus Eusparassus and notes on the RTA-clade and “Laterigradae.”Molecular Phylogenetics and Evolution, 74, 48–65. http://doi.org/10.1016/j.ympev.2014.01.021

Nguyen L. T., Schmidt H. A., Von Haeseler A., & Minh B. Q. (2015). IQ-TREE: A fast and effective stochastic algorithm for estimating maximum-likelihood phylogenies. Molecular Biology and Evolution, 32(1), 268–274. http://doi.org/10.1093/molbev/msu300

Oswald J. A., Overcast I., Mauck W. M., Andersen M. J., & Smith B. T. (2017). Isolation with asymmetric gene flow during the nonsynchronous divergence of dry forest birds. Molecular Ecology, 26(5), 1386–1400. http://doi.org/10.1111/mec.14013

Peres E. A., Sobral-Souza T., Perez M. F., Bonatelli I. A. S., Silva D. P., Silva M. J., & Solferini V. N. (2015). Pleistocene niche stability and lineage diversification in the subtropical spider Araneus omnicolor (Araneidae). PLoS ONE, 10(4). http://doi.org/10.1371/journal.pone.0121543

Posada D. (2008). jModelTest: Phylogenetic model averaging. Molecular Biology and Evolution, 25(7), 1253–1256. http://doi.org/10.1093/molbev/msn083

Pritchard J. K., Stephens M., & Donnelly P. (2000). Inference of Population Structure Using Multilocus Genotype Data. Genetics, 155(2).

Prosdocimi F., Bittencourt D., da Silva F. R., Kirst M., Motta P. C., & Rech E. L. (2011). Spinning Gland Transcriptomics from Two Main Clades of Spiders (Order: Araneae) - Insights on Their Molecular, Anatomical and Behavioral Evolution. PLOS ONE, 6(6). http://doi.org/10.1371/journal.pone.0021634

Ribas C. C., Moyle R. G., Miyaki C. Y., & Cracraft J. (2007).The assembly of montane biotas: linking Andean tectonics and climatic oscillations to independent regimes of diversification in Pionus parrots. Proceedings. Biological Sciences, 274(1624), 2399–408. http://doi.org/10.1098/rspb.2007.0613

Rull V. (2011). Neotropical biodiversity: Timing and potential drivers. Trends in Ecology and Evolution, 26(10), 508–513. http://doi.org/10.1016/j.tree.2011.05.011

Salazar C., Baxter S. W., Pardo-Diaz C., Wu G., Surridge A., Linares M., & Jiggins C. D. (2010). Genetic Evidence for Hybrid Trait Speciation in Heliconius Butterflies. PLOS Genetics, 6(4), e1000930. Retrieved from https://doi.org/10.1371/journal.pgen.1000930

Simon C., Frati F., Beckenbach A., Crespi B., Liu H., & Flook P. (1994). Evolution, Weighting, and Phylogenetic Utility of Mitochondrial Gene Sequences and a Compilation of Conserved Polymerase Chain Reaction Primers. Annals of the Entomological Society of America, 87(6), 651–701. http://doi.org/10.1093/aesa/87.6.651

Smith B. T., McCormack J. E., Cuervo A. M., Hickerson M. J., Aleixo A., Cadena C. D., & Brumfield R. T. (2014). The drivers of tropical speciation. Nature, 515(7527), 406–9. http://doi.org/10.1038/nature13687

Smouse P. E., Long J. C., & Sokal R. R. (1986). Multiple Regression and Correlation Extensions of the Mantel Test of Matrix Correspondence. Systematic Zoology, 35(4), 627. http://doi.org/10.2307/2413122

Tamura K., Stecher G., Peterson D., Filipski A., & Kumar S. (2013). MEGA6: Molecular evolutionary genetics analysis version 6.0. Molecular Biology and Evolution, 30(12), 2725–2729. http://doi.org/10.1093/molbev/mst197

Teixeira M., Prates I., Nisa C., Silva-Martins N. S. C., Strüssmann C., & Rodrigues M. T. (2016). Molecular data reveal spatial and temporal patterns of diversification and a cryptic new species of lowland Stenocercus Duméril & Bibron, 1837 (Squamata: Tropiduridae). Molecular Phylogenetics and Evolution, 94(Part A), 410–423. http://doi.org/https://doi.org/10.1016/j.ympev.2015.09.010

Thompson J. D., Higgins D. G., & Gibson T. J. (1994). CLUSTAL W: improving the sensitivity of progressive multiple sequence alignment through sequence weighting, position-specific gap penalties and weight matrix choice. Nucleic Acids Research, 22(22), 4673–4680. http://doi.org/10.1093/nar/22.22.4673

Toews D. P. L., & Brelsford A. (2012). The biogeography of mitochondrial and nuclear discordance in animals. Molecular Ecology, 21(16), 3907–3930. http://doi.org/10.1111/j.1365-294X.2012.05664.x

Turchetto-Zolet A. C., Pinheiro F., Salgueiro F., & Palma-Silva C. (2013). Phylogeographical patterns shed light on evolutionary process in South America. Molecular Ecology, 22(5), 1193–1213. http://doi.org/10.1111/mec.12164

Werneck F. P., Gamble T., Colli G. R., Rodrigues M. T., & SitesJrJ. W. (2012). Deep diversification and long-term persistence in the south american “dry diagonal”: integrating continent-wide phylogeography and distribution modeling of geckos. Evolution, 66(10), 3014–3034. http://doi.org/10.1111/j.1558-5646.2012.01682.x

White T. J., Bruns T., Lee S., & Taylor J. (1990). Amplification and Direct Sequencing of Fungal Ribosomal Rna Genes for Phylogenetics. In PCR Protocols: A Guide to Methods and Applications (pp. 315–322). http://doi.org/http://dx.doi.org/10.1016/B978-0-12-372180-8.50042-1

Wittkopp P. J., & Beldade P. (2009). Development and evolution of insect pigmentation: Genetic mechanisms and the potential consequences of pleiotropy. Seminars in Cell & Developmental Biology, 20(1), 65–71. http://doi.org/https://doi.org/10.1016/j.semcdb.2008.10.002

